# Seasonality of food availability influences dietary patterns in two farming districts of Malawi

**DOI:** 10.1101/607606

**Authors:** Tiyezge Ruth Zimba, Agnes Mbachi Mwangwela, Beatrice Mtimuni, Prisca Waluza Chisala

## 1.0 Introduction

In most rural areas Africa, farm households rely on their own food production or local open markets for food. Seasonal variations in food availability are characteristic in such areas due to poor food distribution/system infrastructure. Such variability in food availability sometimes results in a food and health crisis that ruins and kills Africans every year, but its severity and duration vary across households and over time [1]. The situation is exacerbated by the rain-fed farming systems, where smallholder farmers depend on a single rainy season for most of their staple food needs. Rain-fed agriculture is dependent on the unpredictable behavior of the weather conditions in the context of climate change [1]. Poor rain performance, directly affects household food and livelihood security, because it affects yield, which in turn is linked to household food consumption and household cash income [2].

In Malawi, food shortages tend to be seasonal, mainly because the vast majority (>84%) of the smallholder farmers depend on rain-fed agriculture and have small land holding sizes (average of < 1.12 hectare) [3]. Seasonal variations in food prices characterized by lowest grain prices around harvest time and steady rise through the dry season until the next harvest also creates vulnerability for the 42% and 82% rural and urban poor Malawians respectively who rely on buying their food from markets [4]. Rural poor farmers are the most vulnerable to seasonal variations in food availability [5]. Many of these famers depend on agriculture for subsistence and income and have limited access to land, financial resources, and farm inputs [1].

Food availability and accessibility have been reported to affect diet pattern. A study in Texas explored social and environmental influences on children’s diets using focus group discussions and they found that availability, accessibility, television, peer and parent influences influenced consumption of fruits, juices, vegetables and low fat foods [6]. Another study which aimed to evaluate the relationship between the home food environment and Hispanic children’s diet quality, found that home food availability, parental diet and family eating habits were associated with the diet quality of Hispanic children [7].

Seasonal food shortages are a common cause of malnutrition among infants [8]. Anecdote evidence showed that the admission pattern of undernourished children to nutrition rehabilitation units in Malawi followed the trend of food scarcity. Most admissions of severely undernourished children to nutrition rehabilitation units in Malawi occurs during the months of January to February, and similarly the highest number of children are admitted to the supplementary feeding or outpatient therapeutic programs during these months [9].

Adequate nutrition is critical for optimal growth, health and cognitive development of infants. Complementary feeding which starts when breast milk alone is no longer sufficient to meet the nutritional requirements of infants is essential from 6 to 23 months of age. However, complementary feeding in rural areas is highly affected by seasonality of food supplies. Seasonal availability and access to different foods were identified among constraints to successful infant and young child feeding (IYCF) interventions by WHO/UNICEF [8]. To address malnutrition, household guidance on recipes based on seasonal food availability has proved to be essential [10].

In Malawi, there is limited dietary diversification due to lack of understanding of food values, poor choices and feeding practices [11]. High incidence of nutrition-related diseases in infants occur during the critical weaning period between 6 months and one year of age in rural areas of Malawi, attributed to inappropriate infant weaning practices [12].

Child undernutrition is one of the big challenges in Malawi and 37% of the under-five children are stunted [13]. The high stunting levels for under-five children in Malawi led to the Malawi government to put in place a number of strategies to combat child malnutrition. One of such strategies is improving women’s nutrition and care before, during and after pregnancy and ensure the consumption of a diversified diet made with foods from the six food groups. This study aimed to determine the seasonal food availability patterns in Dedza and Balaka for the development of seasonal food availability calendar (SFAC) as one of the nutrition interventions to reduce stunting in Malawi. SFAC is one of the tools that can be used to raise awareness to recurring food shortages and helps in developing seasonal complementary food recipes [8].

The study also aimed to determine seasonal variation in the dietary patterns of pregnant women, lactating women and children aged between 12 and 23 months. The first 1000 days of human life, from conception to the age of 2 years, are a critical stage when vital organs such as the brain and its interconnections with the rest of the body are formed and there is rapid growth and development [14]. Extended periods of insufficient nutrient intake for a child during this period can result in permanent damage through stunting [15]. Therefore, the study drew associations between seasonal food availability and dietary patterns of pregnant women, lactating women and children aged between 12 and 23 months.

## 2.0 Materials and Methods

### 2.1 Study area and seasons

The study was conducted in Dedza and Balaka districts, Central and Southern Malawi respectively (Figure 1) during March to December 2015. Dedza has 10 Extension Planning Areas (EPAs) with 169 sections and 197, 492 farming families and Balaka has 6 EPAs consisting of 83 sections with 125, 444 farming families (Ministry of Agriculture and Food Security, 2013). Data were collected in four quarters (March, June, September and December). The seasons in this study were divided into three following Malawi meteorology department description i.e. Warm wet season (November to April), cool dry winter season (May to August) and hot dry season (September to October) [16].

**Figure 1:**
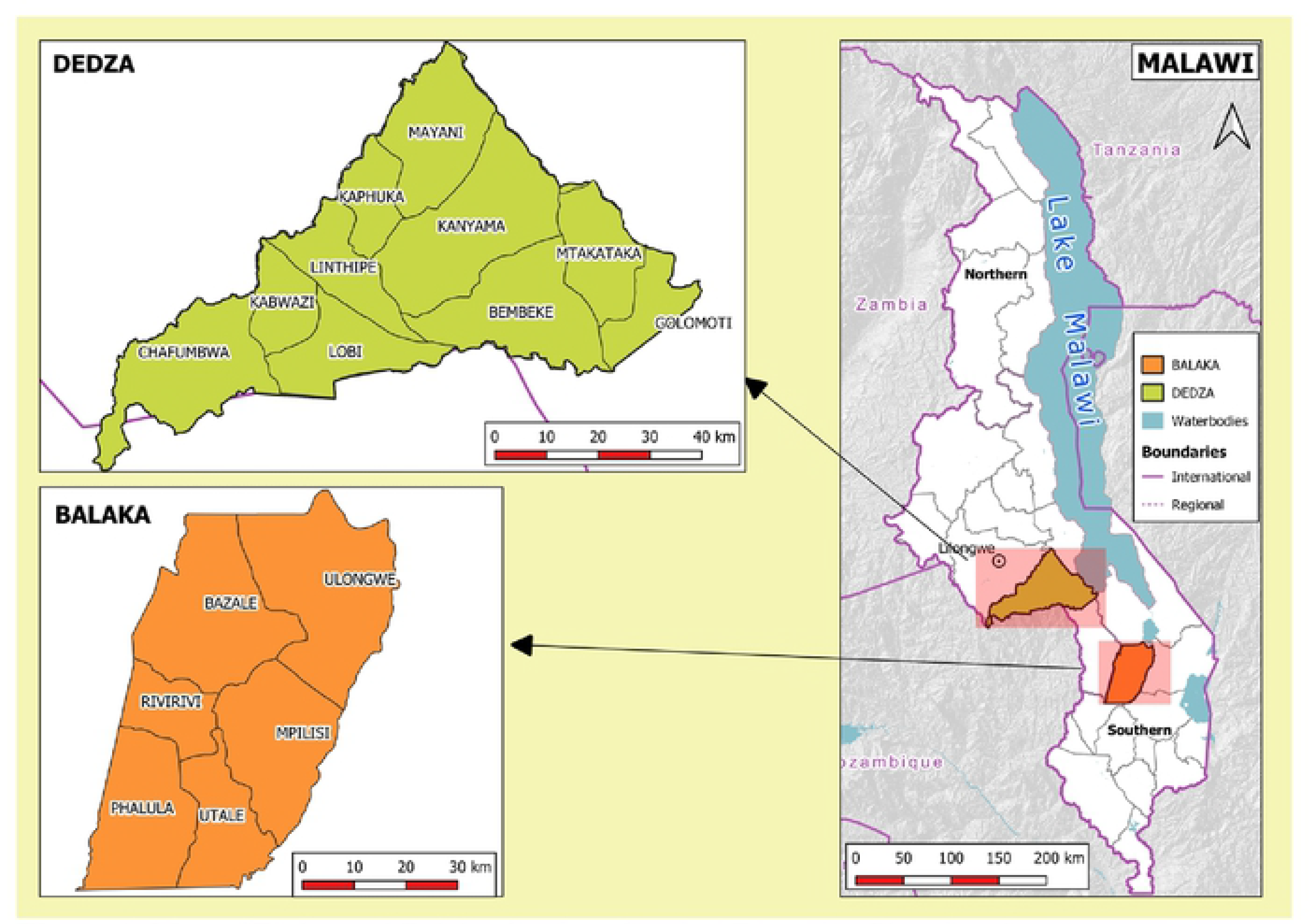
Map of Dedza and Balaka districts showing Extension Planning Areas

### 2.2 Study design and study population

The study was cross sectional employing both qualitative and quantitative data collection methods. The population comprised of pregnant or lactating women and children aged between 12 and 23 months living in 10 EPAs of Dedza and 6 EPAs of Balaka during March to December 2015. The study protocol was reviewed and approved by the Faculty of Food and Human Sciences at LUANAR. Permission to conduct the study was sought from Dedza and Balaka District Commissioner and from Dedza and Balaka District Agricultural Development Officers. Written and thumb-printed informed consent was also obtained from the participants.

### 2.3 Sampling of participants and data collection methods

#### Household survey

The study population was separated into mutually exclusive, homogeneous Extension Planning Areas (EPAs), thus 10 EPAs in Dedza and 6 EPAs in Balaka. A two stage stratified sampling technique was used. First proportional probability sampling was used to randomly sample five EPAs from the two districts that is; three EPAs (Kabwazi, Linthipe and Bembeke) in Dedza and two (Mpilisi and Bazale) EPAs in Balaka (Figure 1). Then purposive sampling technique was used to sample households. Sample size of 160 households with pregnant or lactating women and or children aged between 12 and 23 months was used targeting 32 households in each of the 5 EPAs.

Household survey was used to collected information on dietary patterns and household seasonal food availability. The interviewees included pregnant or, lactating women and mothers of children aged between 12 and 23 months. Due to limited number of children who were eligible throughout the study, all children who were at least within the eligible age group were recruited. A participant was replaced if a household had moved from the area or if the child was no longer within the eligible age range. In the case where a registered pregnant woman gave birth, they were moved to the lactating group. A Semi-structured questionnaire was used to collect data on seasonal availability of foods and a 179 item Food frequency questionnaire validated by a food record was used to determine the dietary patterns. The reference time period for the FFQ was a month and the response categories included; daily, weekly or monthly consumption of food items.

#### Focus group discussions

The participants were selected purposively by targeting 6 to 15 women who were either pregnant, lactating or had children aged between 12 and 23 months from different villages within an EPA in all the 16 EPAs. Focus group discussion participants were not involved in the household survey and data was collected from the same participants in all the four quarters. The discussions were guided by a checklist to collect information on available food with the aid of audio voice recorders.

### 2.4 Data analysis

Principal component analysis (PCA) was used to identify the diet patterns captured using a 179 item FFQ and the data was analyzed using SPSS Version 20. Applicability of factor analysis was verified by Kaiser-Meyer-Olkin (KMO) measurement of adequacy. The presence of correlations between food groups was tested using the Bartlett test of sphericity and was accepted when it was significant at p<0.05. Varimax rotation was used to extract the factors, Eigen value of >1.0 was used to decide the number of components to return and total variance explained indicated how much of the variability in the data was modeled by the extracted factors.

The audio data captured by an audio recorder was transcribed verbatim into written words manually, using Microsoft word. The typed interview transcripts were laid out on left half of the page while keeping the right half margin for writing the codes and notes. The transcript was made anonymous by tagging each voice (e.g. voice A, voice B) to distinguish multiple voices. Coding was done by making notes in the right margin adjacent to each line paragraph. Apart from that, significant participant’s quotes worthy of attention were highlighted. The codes from all the transcripts were drawn together to come up with themes.

Association between food availability and diet pattern were derived by dividing total number of all foods in the diet with total number of food which were shown to be adequate or scarce and multiplying that number with one hundred. Associations then were drawn by looking at the percentage level of availability of the foods that were in the diet.

## 3.0 Results and Discussions

Most of the participants were young married mothers, who had dropped out of primary school and earned their income through farming (Table 1). Only 31% of girls in Malawi complete primary school and 11% graduate from secondary school, such that one in three of the Malawian girls marry and/or experience at least one pregnancy before her 20^th^ birthday [18]. Maternal education has often been associated with positive child health and nutritional outcomes. In Haiti, maternal education was associated with greater dietary diversity in young children’s diets [19]. Considering that 64.5% of the participants in this study did not finish primary school, this may likely affect the way they accept and adhere to nutrition messages.

**Table 1:**
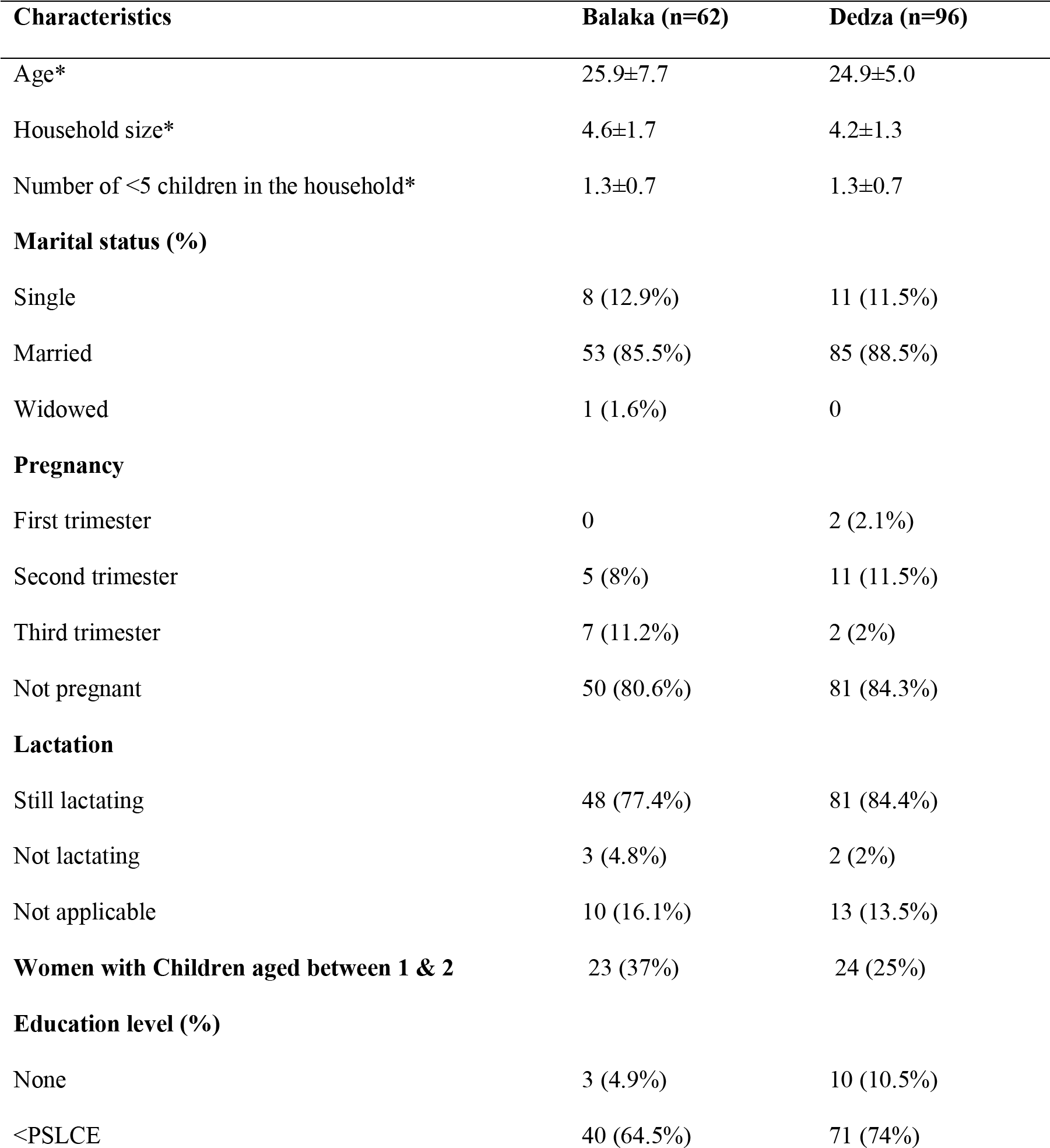
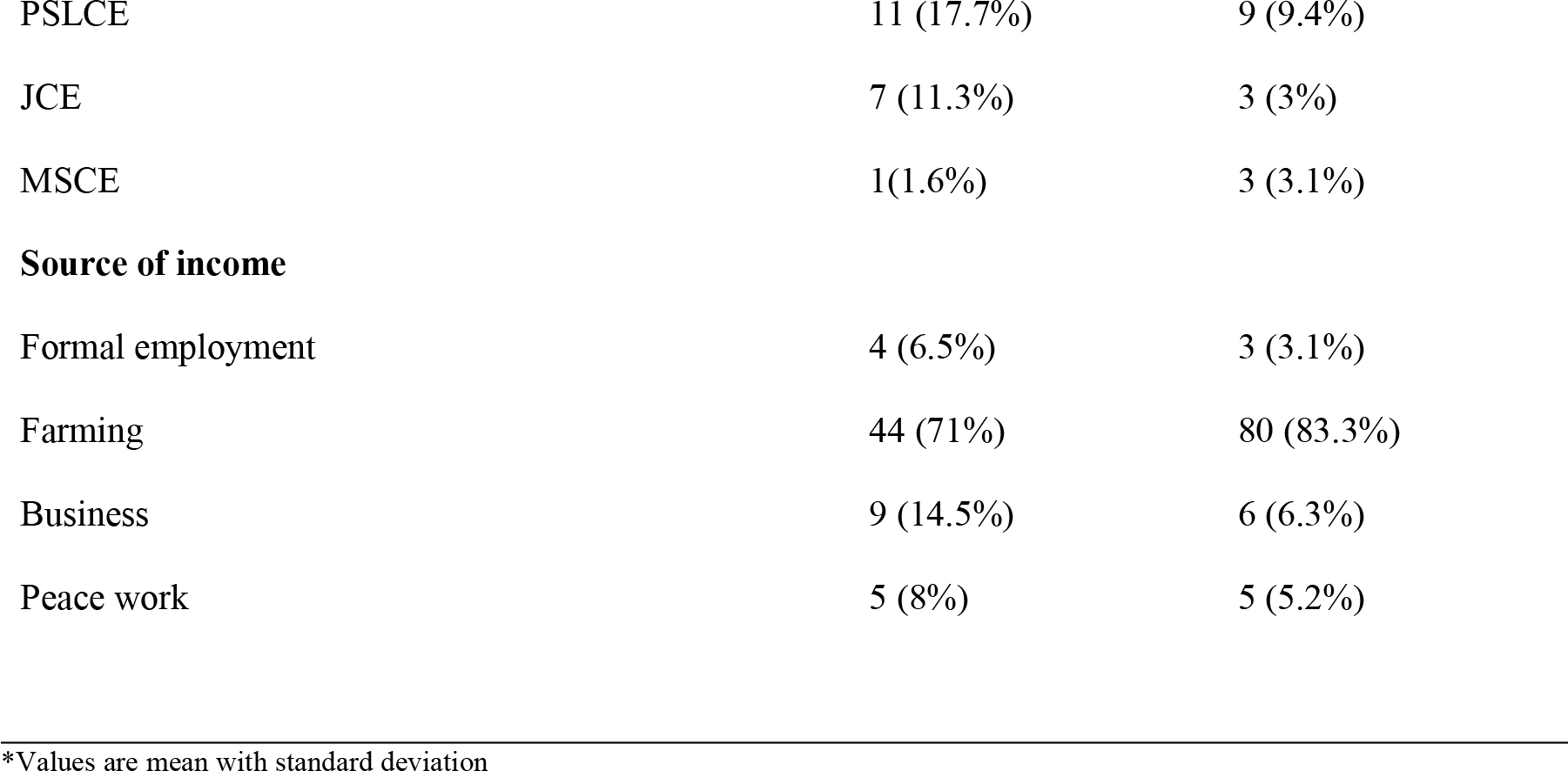
Demographic characteristics of household survey respondents

The cool dry winter season had more variety of foods available as compared to the other seasons. This is because the cool dry winter season coincides with the crop harvest period, and food is readily available through own farms production. However, when the stocks run dry, the households become dependent on purchase [4]. On the other hand, daily meal frequency reduced in cool dry winter season from 3 to 2 meals per day in the hot dry season and 1 meal per day in warm wet season respectively. Similar meal frequencies were reported in Lilongwe district of Malawi, where meal frequency was higher (3 meals per day) immediately after harvest than during pre-harvest period (1 meal per day) [20]. Similarly in Mwense district of Zambia, meal frequency for infants reduced from four to one or two meal per day with change in season [8].

In both districts, maize was the main staple food and it was adequately available all year round (Table 2). This is because maize is a dominant staple food for Malawi supplying more than 54% of the food calories [21]. Common beans were the most common legume in both districts, and were mostly grown in Dedza, while Balaka was a net importer. Legumes grown in Dedza and Balaka were for subsistence and cash crop, however priority was for cash. Economic necessities rather than own consumption preference influence cultivation and sale of food crops. Households sale portions of their harvest in order to buy chemical fertilizers and to meet costs for clothing, schooling, medical services, transport and maize milling” [22]. Vegetables were least available during the hot dry season. In a trials of improved practice (TIPs) study, Malawian mothers raised a general concern that lack of vegetables in the dry season was one of the reasons that made it difficult to adhere to counseling messages for child feeding [8]. In order to improve year-round availability of vegetables, most of the households preserved leafy vegetables through direct sun drying. In times of plenty, vegetables are made in part imperishable by blanching and drying and are stored in traditional storage bags called “*zikwatu*” for future use [22] [24]. Unfortunately, sun drying exposes the food directly to the sun, compromising nutrient retention and hygiene as opposed to solar drying [23]. On the other hand indigenous vegetables were more adequately available in the warm wet season. Another study supports, that indigenous vegetables are abundant during the rainy season, where they grow around family homesteads [24].

**Table 2:** Foods available in Balaka and Dedza districts at different seasons

| Name of food | District | Seasons |  |  |  |  |  |  |  |  |  |  |  |
| --- | --- | --- | --- | --- | --- | --- | --- | --- | --- | --- | --- | --- | --- |
|  |  | Warm wet |  |  |  |  |  | Cool dry winter |  |  |  | Hot dry |  |
|  |  | N | D | J | F | M | A | M | J | J | A | S | O |
| STAPLES |  |  |  |  |  |  |  |  |  |  |  |  |  |
| Maize<br>(*Zea mays*) | Balaka | M | M | M | FM | FM | FM | FM | FM | FM | FM | M | M |
|  | Dedza | FM | FM | M | M | FM | FM | FM | FM | FM | FM | FM | FM |
| Irish potatoes<br>(*Solanum tuberosum*) | Balaka | M | M | M | M | M | M | M | M | M | M | M | M |
|  | Dedza | M | M | M | M | M | FM | FM | FM | FM | FM | FM | FM |
|  |  | Warm wet |  |  |  |  |  | Cool dry winter |  |  |  | Hot dry |  |
|  |  | N | D | J | F | M | A | M | J | J | A | S | O |
| Sweet potatoes<br>(*Ipomoea batatas*) | Balaka | M | M |  |  | FM | FM | FM | FM | FM | FM | M | M |
|  | Dedza |  |  |  |  | FM | FM | FM | FM | FM | FM | FM |  |
| Cassava<br>(*Manihot esculenta*) | Balaka | M | M | M | M | M | FM | FM | FM | FM | FM | M | M |
|  | Dedza | FM | FM | FM | FM | FM |  |  |  |  | FM |  |  |
| Rice<br>(*Oryza sativa*) | Balaka | M | M | M | M | M | M | M | M | M | M | M | M |
|  | Dedza | M | M | M | M | M | M | M | M | M | M | M | M |
| LEGUMES |  |  |  |  |  |  |  |  |  |  |  |  |  |
| Soya beans<br>(*Glycine max*) | Balaka |  |  |  |  | FM | FM | FM | FM | FM | FM |  |  |
|  | Dedza | M | M | M | M | M | F | F | F | FM | FM | FM | M |
| Dry pigeon peas<br>(*Cajanus cajan*) | Balaka | FM | FM | FM | FM | FM | F | F | FM | FM | FM | FM | FM |
|  | Dedza |  |  |  |  |  |  |  |  |  |  |  |  |
| Dry common bean<br>(*Phaseolus spp*) | Balaka | M | M | M | M | M | M | M | M | M | M | M | M |
|  | Dedza | FM | M | M | M | F | F | FM | FM | FM | FM | FM | FM |
| VEGETABLES |  |  |  |  |  |  |  |  |  |  |  |  |  |
| Cabbage<br>(*Brassica oleracea var. capitata*) | Balaka | M | M | M | M | M | M | M | M | M | M | M | M |
|  | Dedza | M | M | M | M | M | FM | FM | FM | FM | M | M | M |
| Rape<br>(*Brassica napus var. oleifera*) | Balaka | M | M | M | M | M | FM | FM | FM | FM | FM | M | M |
|  | Dedza | M | M | M | M | M | FM | FM | FM | FM | FM | M | M |
| Mustard Indian<br>(*Brassica juncea*) | Balaka | M | M | M | M | M | M | M | M | M | M | M | M |
|  | Dedza | M | M | M | M | M | FM | FM | FM | FM | FM | M | M |
| Sweet potato leaves<br>(*Ipomoea batatas*) | Balaka | F | F | F | F | F | F | F | F | F | F | F | F |
|  | Dedza | F | F | F | F | F | F | F | F | F | F | F | F |
| Pumpkin leaves<br>(*Cucurbita species*) | Balaka | M | M | F |  | F | F | F | FM | FM | FM | M | M |
|  | Dedza | FM | FM | F | F | F | F | F | FM | FM | FM | FM | FM |
|  |  | Warm wet |  |  |  |  |  | Cool dry winter |  |  |  | Hot dry |  |
|  |  | N | D | J | F | M | A | M | J | J | A | S | O |
| Cassava leaves<br>(*Manihot esculenta*) | Balaka | F | F | F | F | F | F | F | F | F | F | F | F |
|  | Dedza | F | F | F | F | F | F | F | F | F | F | F | F |
| Amaranth<br>(*Amaranthus hybridus*) | Balaka | M | F | F | F | F | F | FM | FM | FM | FM |  | M |
|  | Dedza | F | F | M | F | F | FM | FM | FM | M | M | M | M |
| Blackjack<br>(*Bidens pilosa*) | Balaka |  | F | F | F | F | F | F | F | F |  |  |  |
|  | Dedza | M | M | M | M | M | M | M | M | FM | FM | FM | FM |
| Ye-be<br>(*Cardeauxia edulis*) | Balaka |  | F | F | F | F | F | F | F |  |  |  |  |
|  | Dedza |  | F | F | F | F | FM | FM |  |  |  |  |  |
| Sweet corn<br>(*Zea mays*) | Balaka | M | M |  | F | F | F | FM | FM | M | M | M | M |
|  | Dedza | M | M |  | F | F | F | F | FM | FM | FM | M | M |
| Tomato<br>(*Lycopersican esculentum*) | Balaka | M | M | M | M | M | M | M | M | M | M | M | M |
|  | Dedza | M | M | M | M | M | M | M | M | M | M | M | M |
| Okra<br>(*Hibiscus esculentus*) | Balaka | M | M |  | M | FM | F | F | F | FM | FM |  | M |
|  | Dedza | M | M | M | FM | FM | FM | FM | FM | FM | M | M | M |
| Onion<br>(*Allium cepa*) | Balaka | M | M | M | M | M | M | M | M | M | M | M | M |
|  | Dedza | M | M | M | M | M | M | M | M | M | M | M | M |
| ANIMAL PRODUCTS |  |  |  |  |  |  |  |  |  |  |  |  |  |
| Usipa/Bonya fish<br>(*Engraulicypris sardella-Large*) | Balaka | M | M | M | M | M | M | M | M | M | M | M | M |
|  | Dedza | M | M | M | M | M | M | M | M | M | M | M | M |
| Cat fish-Mlamba/Anjolinjo<br>(*Clarias gariepinus*) | Balaka | M | M | M | M | M | M | M | M | M | M | M | M |
|  | Dedza | M | M | M | M | M | M | M | M | M | M | M | M |
| Utaka<br>(*Copadichromis, Otopharynx*) | Balaka | M | M | M | M | M | M | M | M | M | M | M | M |
|  | Dedza | M | M | M | M | M | M | M | M | M | M | M | M |
| Micheni | Balaka |  |  | M | M | M | M | M | M | M | M | M | M |
|  |  | Warm wet |  |  |  |  |  | Cool dry winter |  |  |  | Hot dry |  |
|  |  | N | D | J | F | M | A | M | J | J | A | S | O |
| *(Rhamphochromis sp.)* | Dedza | M | M | M | M | M | M | M | M | M | M | M | M |
| *Matemba*<br>*(Barbus paludinosus)* | Balaka | M | M | M | M | M | M | M | M | M | M | M | M |
|  | Dedza | M | M | M | M | M | M | M | M | M | M | M | M |
| Chicken | Balaka | FM | FM | FM | FM | FM | FM | FM | FM | FM | FM | FM | FM |
|  | Dedza | FM | FM | FM | FM | FM | FM | FM | FM | FM | FM | FM | FM |
| Eggs | Balaka | FM | FM | FM | FM | FM | FM | FM | FM | FM | FM | FM | FM |
|  | Dedza | FM | FM | FM | FM | FM | FM | FM | FM | FM | FM | FM | FM |
| Pork | Balaka | FM | FM | FM | FM | FM | FM | FM | FM | FM | FM | FM | FM |
|  | Dedza | FM | FM | FM | FM | FM | FM | FM | FM | FM | FM | FM | FM |
| Goat Meat | Balaka | FM | FM | FM | FM | FM | FM | FM | FM | FM | FM | FM | FM |
|  | Dedza | FM | FM | FM | FM | FM | FM | FM | FM | FM | FM | FM | FM |
| Beef | Balaka |  |  | M | M | M | M | M | M | M | M |  |  |
|  | Dedza |  |  | M | M | M | M | M | M | M | M |  |  |
| Cow milk | Balaka |  |  | M |  |  | M | M | M | M | M |  |  |
|  | Dedza | M | M | M | M | M | M | M | M | M | M | M | M |
| FRUITS |  |  |  |  |  |  |  |  |  |  |  |  |  |
| Jujube<br>*(Ziziphus mauritiana)* | Balaka |  |  |  |  | FM | FM |  | FM | FM | FM | FM |  |
|  | Dedza |  |  |  |  |  | FM | FM | FM | FM | FM |  | FM |
| Baobab<br>*(Adansonia digitata)* | Balaka |  |  |  |  | FM | FM | FM | FM | FM |  |  |  |
|  | Dedza |  |  |  |  | FM | FM | FM |  |  |  |  |  |
| Ripe Bananas *(Musa paradisiaca)* | Balaka | M | M | FM | FM | FM | FM | M | M | M | M | M |  |
|  | Dedza | M | M | FM | FM | FM | FM | FM | FM | FM | FM | M |  |
| Mangoes *(Mangifera indica)* | Balaka | F | F | FM | FM |  |  |  |  |  | FM | F |  |
|  | Dedza | F | F | F |  | FM |  |  |  |  | FM |  |  |
|  |  | Warm wet |  |  |  |  |  | Cool dry winter |  |  |  | Hot dry |  |
|  |  | N | D | J | F | M | A | M | J | J | A | S | O |
| Guava (*Psidium guajava*) | Balaka |  |  |  | FM | FM | FM | FM | FM |  |  |  |  |
|  | Dedza |  |  |  | FM | FM | FM | FM | FM | FM |  |  |  |
| OIL FOODS |  |  |  |  |  |  |  |  |  |  |  |  |  |
| Vegetable Cooking oil | Balaka | M | M | M | M | M | M | M | M | M | M | M | M |
|  | Dedza | M | M | M | M | M | M | M | M | M | M | M | M |
| Soya beans (*Glycine max*) | Balaka |  |  |  |  | FM | FM | FM | FM | FM | FM |  |  |
|  | Dedza | M | M | M | M | M | F | F | F | FM | FM | FM | M |
| Dry groundnuts (*Arachis hypogaea*) | Balaka | M | M | M | M | M | FM | FM | FM | FM | FM | M | M |
|  | Dedza | FM | M | M | M | M | F | F | F | FM | FM | FM | FM |
| Avocado pears (*Persea americana*) | Balaka |  |  |  | M | M | M | M | M | M | M |  |  |
|  | Dedza |  |  |  |  | M | M | M | M | M | M |  |  |

Key: N- November, D- December, J- January, F- February, M-March, A- April, M- May, J-June, J-July, A- August, S- September, O- October
Adequately available Scarcely available Not available
Source of food: (F) individual household farming; (M) purchase from the market; (FM) purchase from the market and individual household farming

Exotic vegetables, animal and oil foods, were predominantly obtained from the market in both districts, therefore consumption of these foods highly depends on financial resources of a household. Fish is the main source of animal protein for poor rural households in Malawi. Forty percent of the total protein supply comes from fish, which provides over 60% of dietary animal protein intake of Malawians [25]. Households involved in animal farming don’t usually consume the animals that they farm, because animal farming has a prestigious value attached to it and most of the households would only consume their own animals during ceremonies. Consumption of fruits was lower than that of vegetables and only half of all households in Malawi consume fruits [26]. A participant in Dedza mentioned that “*In Malawi, households do not usually grow fruits, most of the fruits that are available are naturally available in the communities*”. When it comes to government agricultural programs and extension workers expertise in Malawi, there is also a bias towards staple crops and cereals, with a small focus on horticulture and fruit tree production [27]. On the other hand, vegetable-cooking oil was adequately available across all the seasons in both districts. One of the participants in Balaka mentioned that “*unlike some years back, with the awareness of the importance of inclusion of oil foods in the diet and the preferred taste of fried foods, most of the households now make sure that they cook meals using cooking oil for at least once a week*”. The participants also explained that most households can now afford cooking oil because vendors provide cooking oil in small sachets with a starting price of K50 (around 0.069$) which is cheaper as compared to commercial packaging which starts from a 500ml bottle at MK750 (around 1$).

A Malawian recommended diet comprising of six food groups was not observed among the study participants across all the seasons. A recommended diet for Malawians including pregnant and lactating women as well as children aged between 12 and 23 months, is a diet comprising of food from the six food groups of Malawi; staples, legumes, vegetables, animal products, fruits and oils [17]. Increasing the variety of foods across and within food groups is recommended to ensure adequate intake of essential nutrients that promote good health. However, diets comprising of at least four food groups which is considered acceptable, were observed in the warm wet season and hot dry season; “*vegetable, oil, staple, fish and legume based diet*” observed in Balaka in the warm wet season, “*vegetable, legume, oil, staple and fruit based diet*” observed in Balaka in the hot dry season and “*legume, fish, vegetable and staple based diet*” observed in Dedza in the hot dry season (Table 3). Other studies in rural Malawi have reported that 33% of rural Malawians consumed food from less than five out of the six food groups [4].

**Table 3:**
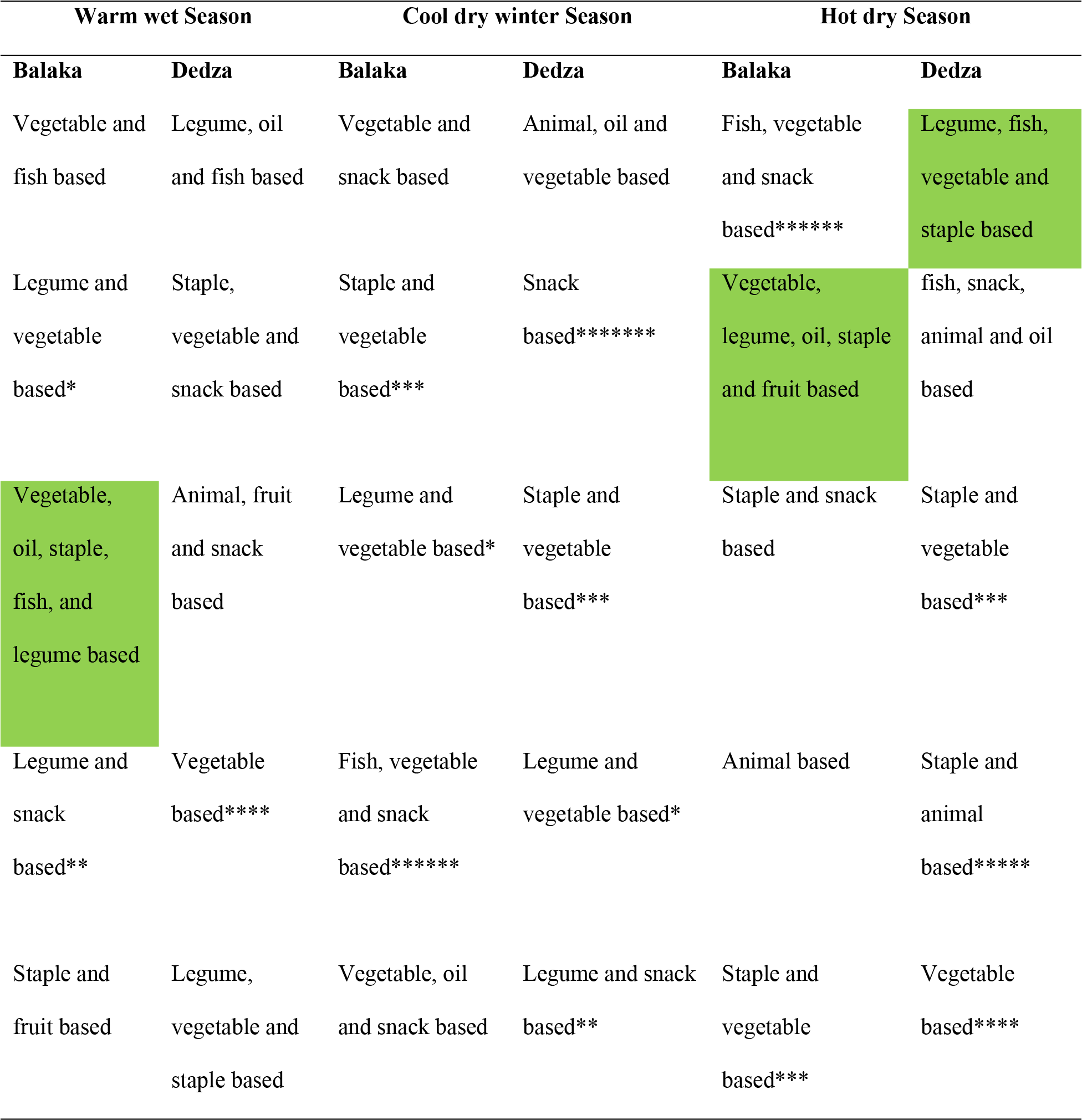
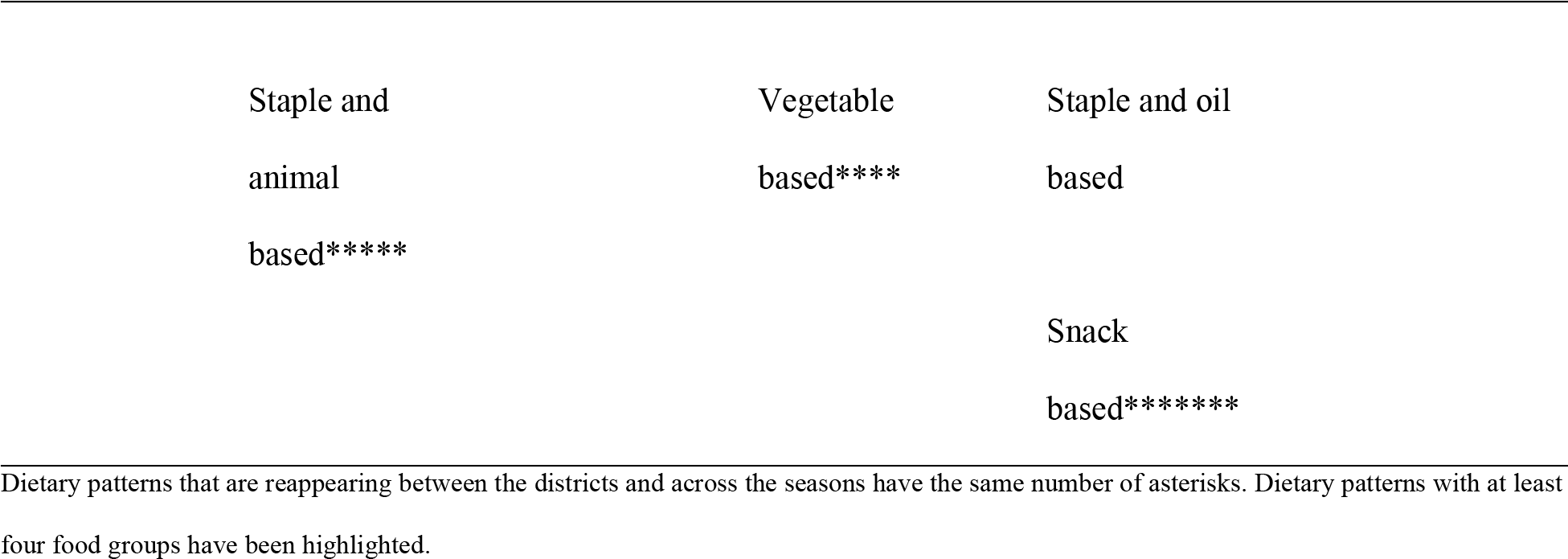
Summary of dietary patterns for pregnant and lactating women and children aged between 6 and 23 months for Dedza and Balaka at different seasons

Diets comprising of food made from at least four food groups were observed in the warm wet season and hot dry season, showing that malnutrition cases are more likely to increase in the cool dry winter season. This is contrary to expectation, since the cool dry season coincides with harvest time, when households had access to a variety of foods as indicated by the seasonal food availability calendars (Table 2). In ability to consume a balanced diet in the cool dry winter season may be due to a number of reasons. Inadequate nutrition knowledge among women and men farmers contributes to unbalanced nutrition in Malawi [22]. On the other hand, people’s attitude towards health and nutrition advice also affects diet that is followed. In one study, 85.6% of the students were familiar with the concept of balance of nutrients in food, but only 7% of them used it in their diets [28]. Time is also a challenge, this being a harvest season, women are usually busy in the fields harvesting, hence they may not have adequate time to prepare nutritious meals. It should also be noted that food choices are largely driven by taste, cost, and convenience [29]. Dietary guidelines tend to emphasize good nutrition, rarely taking food preferences, food prices, or diet costs into account. The ability to follow a healthy diet depends on having sufficient knowledge, money, and time, which low-income families often lack.

A brief overview shows that the constituents of the diets were food items that were adequately available in homes as compared to those that were adequately available at the market. In the warm wet season, 87% and 82% of the food that comprised the diets of Balaka and Dedza respectively were adequately available in the home, while in the cool dry winter season, 92% of the foods in Balaka diets and 74% of the food in the Dedza diets were adequately available. Likewise in the hot dry season 65% and 67% of the foods in the diets of Balaka and Dedza respectively were adequately available (Table 2 and 3). Several studies indicate existence of positive associations between food consumption and availability of foods in the home [30]. One study found that intakes of fruit and vegetables were positively associated with household availability and that consumption of milk increased when milk was available in the home [31]. Yet in another study, it was found that adolescents consumed less snacks and sweetened beverages when these foods were not available in the home [32].

In conclusion, the findings of the present study supported by other studies, has revealed that nutrition knowledge is essential to achieving adequate food utilization. That home food availability is associated with consumption patterns as opposed to market food availability, which is entangled with issues of accessibility. Enhancing the nutrition attitudes, knowledge and practices of people is important because this would lead to more food-conscious society, which could be predisposing factors for improving eating habits and adopting a healthy diet. Simple messages on behavior change mechanisms on improving dietary pattern and provision of different types of recipes containing locally available foods will help households to make better food choices which could be a component of dietary change intervention.

## Acknowledgements

The authors would like to thank Mr. Vincent Mlotha, for his assistance with data analysis, Mr. Clyde Kalima, for his assistance with development of maps and the participants, without whose cooperation this study would not have been a success.

